# deepTools4: Faster, Scalable, and Modernized Genomic Data Analysis

**DOI:** 10.64898/2026.11.11.765524

**Authors:** Ward Deboutte, Adrian Salatino, Wolfgang Maier, Pavankumar Videm, Mohit Navandar, Galina Erikson, Saim Momin, Juan Caballero-Perez, Leily Rabbani, Lucille Lopez-Delisle, Björn Grüning, Thomas Manke

**Affiliations:** Max Planck Institute of Immunobiology and Epigenetics, Freiburg, Germany; Bioinformatics Group, Department of Computer Science, University of Freiburg, Freiburg, Germany; Department of Molecular Medicine and Surgery, Karolinska Institutet, Stockholm, Sweden; Department of Genetic Medicine and Development, Faculty of Medicine, University of Geneva, Geneva, Switzerland; HTW Berlin, School of Computing, Communication and Business, Berlin, Germany

**Keywords:** high-throughput sequencing, ChIP-seq, ATAC-seq, genome-wide analysis, read normalization, sequencing data visualization, quality control, epigenomics, bioinformatics, genome coverage

## Abstract

The deepTools suite is widely used for quality control, summarization, and visualization of high-throughput sequencing data. However, as genomic datasets grow in scale and depth, existing tools often become computational bottlenecks. We present deepTools4, a modernized framework designed for high performance, scalability, and enhanced usability, while maintaining seamless integration with the Galaxy platform. Core functionality has been rewritten and thoroughly benchmarked, achieving substantial computational speed-ups. Visualization tools have been modernized to provide more intuitive, high-quality graphical outputs. In addition, our testing framework has been expanded to facilitate continuous development and ensure robustness. We demonstrate the utility of deepTools4 across diverse epigenomics and transcriptomics datasets, highlighting its ability to summarize genome-wide signals, apply a wide range of common normalization methods, and visualize enrichment patterns. The enhanced deepTools4 framework delivers a faster, more scalable solution for reproducible genome-wide analysis and interpretation across many sequencing modalities.

**Availability:** https://github.com/deeptools/deepTools/

## Introduction

Advances in high-throughput sequencing technologies have transformed genomics research, enabling the generation of vast amounts of short and long reads at unprecedented scale^1,2^. For more than a decade now, the rate at which sequencing data can be generated has consistently outpaced the capacity to analyze and interpret it^3^. This discrepancy is particularly pronounced in epigenomics, where the complexity and availability of different assays requires not only substantial computational resources, but an iterative and nuanced analytical approach to properly extract biological insights^4^. Unlike primary DNA sequencing, epigenomic signals are often characterized by higher levels of noise and larger variability, necessitating extensive validation, quality control, and comparative analysis to ensure that observed results are genuine biological observations^5^. While the bioinformatics community has developed a wide range of algorithms and tools to scale the first steps of (epi)genomics data processing, such as base calling and genome alignment, the full analytical process did not co-evolve and suffers from fragmented tools, scripts and ad hoc solutions that are usually implemented as a series of disjoint, project-specific scripts. This fragmentation not only hinders reproducibility but often results in substantial performance bottlenecks when scaling to the massive datasets and sample sizes now common in the field. In parallel with the increasing size of genomic datasets, there has been growing interest in the bioinformatics field in leveraging Rust^6^ as a language for high-performance and memory-safe tooling, as it addresses the scalability and robustness challenges that come with the large amount of data generated in epigenomics to date. This is exemplified by the Rust-Bio^7^ ecosystem, which provides several libraries for data structures and algorithms commonly used in bioinformatics, and more recent tools such as bigtools^8^ and GRanges^9^. However, these improvements have primarily benefited tools upstream of the genomics data analysis pipeline and researchers still need to perform the full analytical process, which is often iterative and includes quality controls, different normalization approaches, filtering steps, file and sample comparisons, data integration, and visualization^10,11^. For example, the selection of an appropriate normalization strategy or filtering thresholds often requires multiple rounds of visualization and recalculation before conclusions can be drawn. The deepTools suite aims to simplify and standardize this process, by providing a unified framework for a wide range of applications and a set of standalone tools for interactive data explorations^12^. Moreover, modern bioinformatics tools should facilitate not only expert-level analysis, but also provide accessibility for researchers with less technical background and the training community. This requires well-maintained, documented, and user-friendly software that integrates seamlessly into existing workflows and workflow management systems such as Galaxy^13^, Snakemake^14,15^ and Nextflow^16^. To ensure efficiency for many samples and at genome-wide scale, the deepTools suite presents a set of modular tools that operate on highly standardized data formats and utilizes data-parallelism in all its core components. With the latest release of deepTools4, we introduce significant modernizations aimed at improving the performance, usability, and sustainability of long-term software development. By rewriting core functionalities in Rust, deepTools4 achieves a substantial computational speed-up. Additionally, we have modernized the visualization tools and expanded the testing framework to ensure continuous development, long-term maintainability and seamless integration into platforms like Galaxy. These improvements solidify deepTools as a fast, reliable and scalable open software solution for genomics research and training.

## Results

### The deepTools4 ecosystem: a unified framework for epigenomic analysis

The deepTools suite is designed as a modular framework to support the complete analytical workflow of epigenomic data, from alignment files to publication-ready visualizations (Figure 1). The ecosystem is organized in three functional tiers: tools to process alignment files (filtering and normalization), perform quality control, and tools to produce visualizations. We maintain compatibility with established genomic formats, utilizing BAM or CRAM files for aligned reads, bigWig or bedGraph files for signal scores, and BED or GTF files for genomic regions. To meet the demands of modern large-scale datasets, often requiring signal aggregation over tens of thousands of genomic regions, the core computational engine was rewritten in Rust. By leveraging rust-htslib and bigtools, for parsing BAM/CRAM files and writing bigWig files, respectively, deepTools4 now provides high-performance handling of sequencing data. Complementing this backend overhaul, the plotting functionality has been extensively updated to improve visual clarity and flexibility. Throughout these updates, we have ensured that intermediary files and output files remain compatible and consistent with older versions of deepTools, providing a seamless transition for users with established workflows.

**Figure 1.**
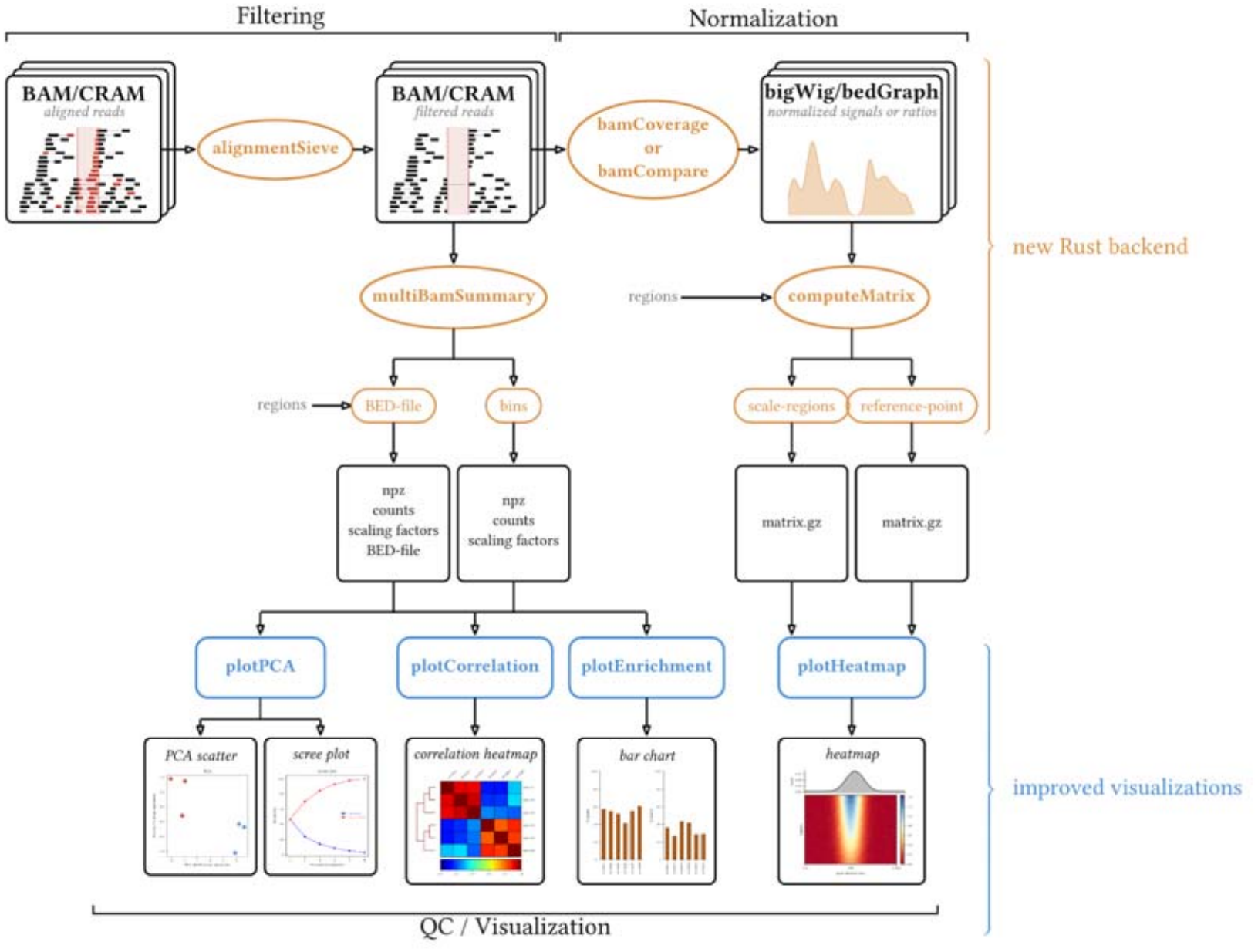
Overview of the rewritten core tools available in deepTools4. The common starting point for most deepTools analyses is a set of BAM or CRAM files with the coordinates of aligned reads for a genome of interest. With a series of efficient tools, these reads can be filtered and manipulated (*alignmentSieve*), binned and normalized for signal extraction (*bamCoverage*), compared for pairwise samples (*bamCompare*), summarized over genomic bins or regions of interest for multiple samples (*multiBamSummary*) or for specific regions of interest (*computeMatrix*). Ultimately, the computed aggregated data can be used to generate standardized output formats and visualizations using several available tools (*plotPCA, plotCorrelation, plotEnrichment, plotHeatmap*).

### Systematic benchmarking shows consistent speed-up over a wide range of conditions

A Snakemake workflow was designed to compare the performance of the rewritten backend. To ensure that the benchmark would reflect both realistic use cases as well as function as a computational stress test, public data from two different organisms, human (*Homo sapiens*) and bread wheat (*Triticum aestivum)* were selected. The human genome serves as the standard benchmark, while the wheat genome represents a substantial challenge due to its large genome size (1.4 x 10^10^ base pairs), hexaploid nature and repetitive genome architecture across 21 chromosomes^17^. For both organisms, whole-genome sequencing, RNA-seq and ChIP-seq datasets were included, to account for differing read densities, distribution patterns and signal profiles (Supplemental table 1). For all five rewritten processing tools, an exhaustive benchmark was set up that spanned up to 20 threads. This allowed us to evaluate the speedup relative to deepTools3 in both low-resource settings (where threads are limited) as well as a high-performance setting to ensure that performance gains are consistent across the entire range. The rewritten backend demonstrates a significant performance increase across the five processing tools over all the threads tested (Table 1). The most dramatic gains were observed with *computeMatrix*, which achieves a near 25-fold reduction in runtime in the low-thread configuration (Figure 2). At 8-16 threads, deepTools4 runs in about 1-13 minutes (median of 4 minutes) versus 5-47 minutes for deepTools3, which corresponds to a 3.5-14-fold speed-up for all tools except for alignmentSieve (Supplemental Figure 1). Peak memory usage increased *bamCoverage, bamCompare* and *computeMatrix*, due to a shift from disk-based intermediate result storage and aggregation in the previous implementation to in-memory buffering in a new shared-memory parallel implementation. Despite this increase, usage remains well within the limits of normal genomic workflows as the peak average memory usage was 8 GB (Table 1, Supplemental figure 2). We conclude that the backend rewrite provides a major efficiency boost for both users with limited local resources and those utilizing high-performance computing clusters.

**Table 1.** Runtime and memory of deepTools4 compared with deepTools3. Per tool, dataset (human and *Triticum aestivum*), and thread count, the speed-up was computed as the mean deepTools3 runtime divided by the mean deepTools4 runtime over five replicates. The geometric mean and median across all datasets and thread counts are denoted in the second and third column, respectively. To summarize the range, the lowest (min) and highest (max) speed-up across all datasets (human and *Triticum aestivum*) were taken at each thread count, and these values were averaged over all thread counts, yielding the avg. min and avg. max values denoted in the third column between brackets. Memory usage is peak resident set size, averaged over all repetition and thread counts for each dataset, and then averaged across datasets.

| **Tool** | **Geometric mean speed-up** | **Median speed-up (avg. min - avg.max)** | **Average memory usage deepTools3 (GB)** | **Average memory usage deepTools4 (GB)** |
|---|---|---|---|---|
| alignmentSieve | 1.89 | 2.19 (1.01 - 4.15) | 1.32 | 0.06 |
| bamCompare | 3.53 | 3.54 (3.26 - 3.95) | 2.12 | 8.00 |
| bamCoverage | 5.37 | 5.14 (4.33 - 7.25) | 1.26 | 3.53 |
| computeMatrix | 10.25 | 10.97 (5.85 - 16.02) | 5.06 | 6.33 |
| multiBamSummary | 5.18 | 5.14 (3.77 - 7.16) | 2.44 | 0.46 |

**Figure 2.**
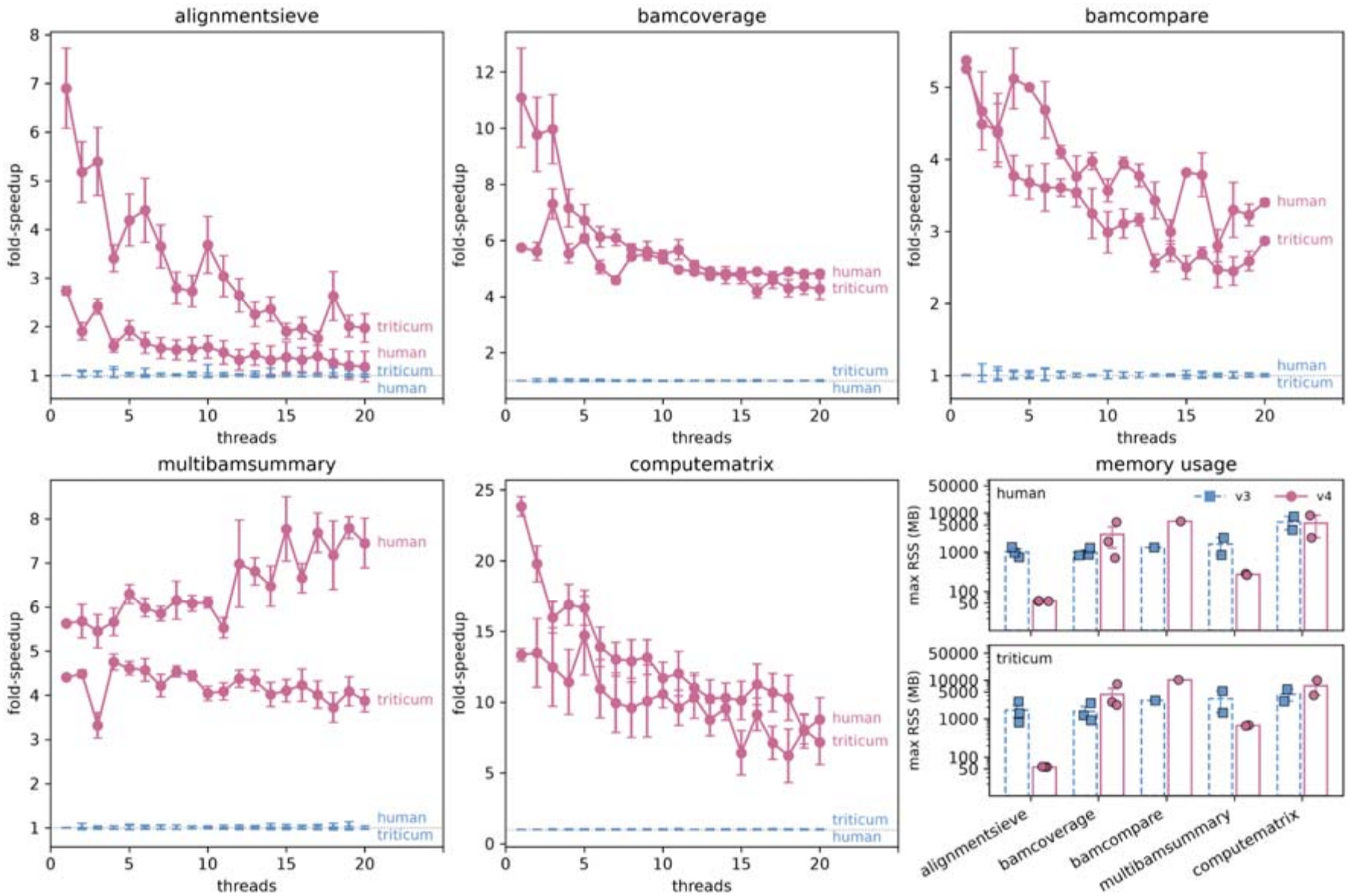
Exhaustive benchmarking reveals a consistent speedup for deepTools4 across a range of threads. Shown is the relative speedup for deepTools4 (circles, purple) compared to deepTools3 (baseline, blue). For deepTools3 only the standard error of the mean is shown. The last panel depicts the maximum memory (maximum resident set size in MB) used by deepTools4 (solid, purple) and deepTools3 (dotted, blue).

### A multi-layered testing framework ensures stability and continued development

Offloading core functionality to Rust necessitated a partial migration of the existing test suite to ensure that the output remains consistent with deepTools3, long-term reproducibility is guaranteed and future development is facilitated. Alongside the new backend, we utilized Rust’s built-in testing framework to implement comprehensive unit tests for all new backend components. Because the Rust rewrite focuses specifically on core functionality (i.e. converting standardized data formats into formats more suitable for visualization and statistical computations), the need for unit tests for the Python codebase and Galaxy integration persists. As of today, the testing framework for deepTools consists of three hierarchical layers. The first encompasses the Rust unit testing framework, which ensures the core functionality works as expected in its native environment. The second layer ensures robustness of the command-line-interface (CLI), visualization functionality and statistical computations, and is based on the pytest framework, as these functions are still implemented in Python. The third layer ensures a seamless integration of deepTools in Galaxy, and includes testing on both the backend level as well as the visualization and computation level (Supplemental table 2). To verify that deepTools4 reproduces the output of deepTools3, we ran both versions on three inputs per organism (human and *Triticum aestivum*) for each tool and compared their outputs value by value (Figure 3). *AlignmentSieve* outputs were identical for all reads in all six samples. For the other tools (*bamCoverage, bamCompare, multiBamSummary and computeMatrix)*, outputs were highly concordant (Pearson correlation > 0.9999) for all runs on an individual basis. Where values differed, the differences were small in absolute terms and negligible relative to the values themselves: relative differences were below 10^-5^ for multiBamSummary and computeMatrix, and below 10^-2^ for nearly all bamCoverage and bamCompare values (Figure 3). The largest absolute differences occurred at loci with extreme read depth (>10^6^ reads per bin), where they correspond to a relative difference of 10^-6^. These differences are primarily attributed to the fact that intermediate computations now use single-precision 32-bit floating points and that deepTools4 writes output with values rounded to two decimal places instead of the six decimal places that are retained in deepTools3, mainly to optimize storage and reduce file sizes.

**Figure 3.**
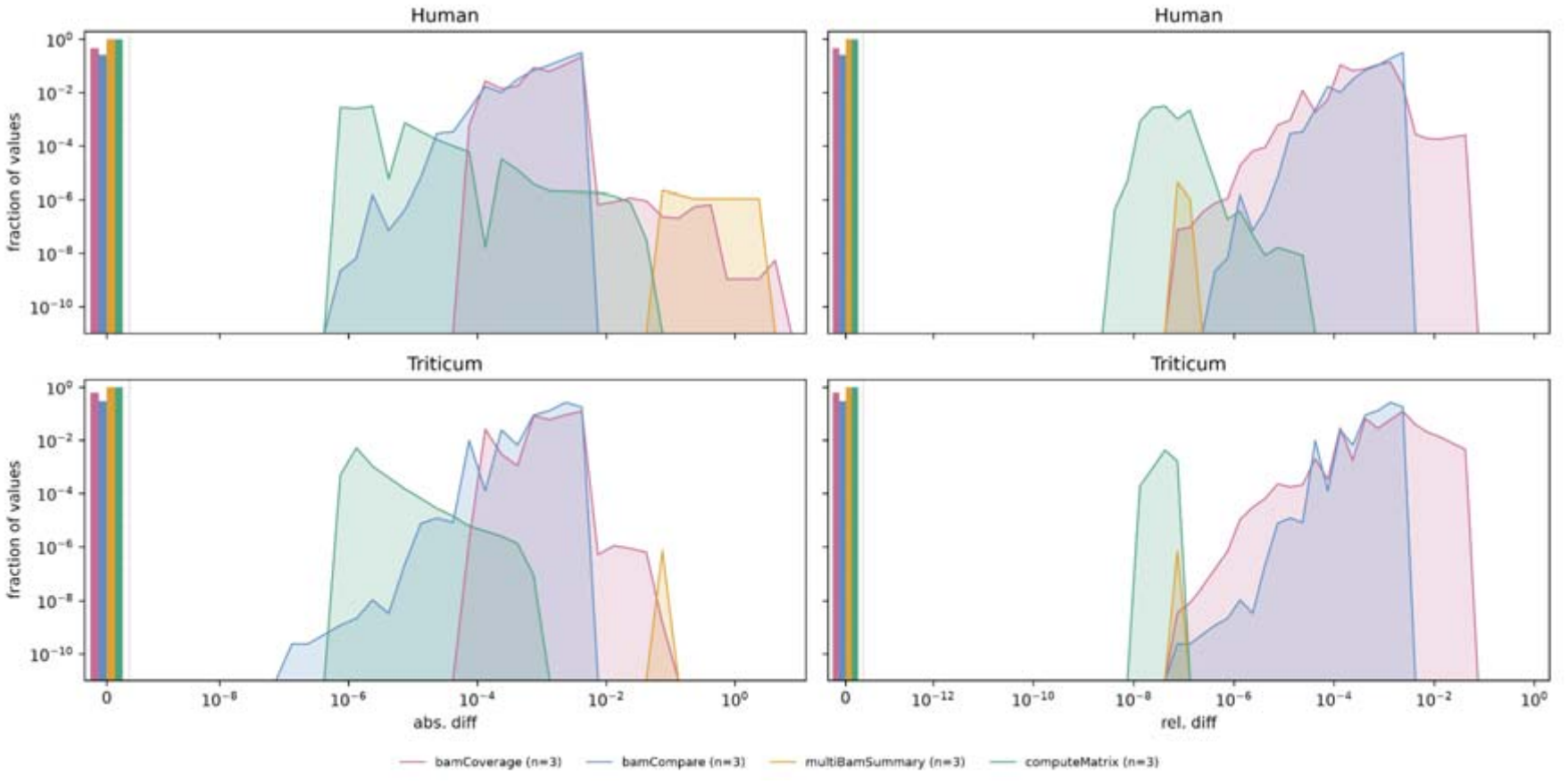
Numerical value comparison of deepTools3 versus deepTools4. Shown are the distributions of absolute (left) and relative (right) differences between deepTools3 and deepTools4 output for human (top) and *Triticum aestivum* (bottom), averaged over three input configurations per tool (bamCoverage: ChIP-seq, RNA-seq and WGS samples, bamCompare: ChIP/WGS, RNA/WGS and ChIP/RNA pairs, computeMatrix and multiBamSummary with 3, 6 and 15 input files where the 6 and 15-file sets were repeats of the ChIP, WGS and RNA samples). Bar plots on the left of the dotted line depict the fraction of identical values (no difference).

### Multimodal integration for the epigenetic characterization of differentially expressed genes

To demonstrate the utility of the deepTools suite in a complex biological context, we applied the framework to a multimodal dataset integrating RNA-seq with seven distinct epigenomic layers. Publicly available data was used to determine differentially expressed genes between MSL2 knock-out (KO) murine neuronal progenitor cells (mNPCs) and wildtype (WT) mNPCs^18^. Sample depth and characteristics were in line with what one would expect for a regular epigenomic dataset (Supplemental table 3). A total of 1,904 differentially expressed genes were detected (894 upregulated in KO, 1,010 downregulated in KO), and from the non-differentially expressed genes 1,000 were selected for downstream visualization, yielding 2,904 genes of interest (Figure 4A). As part of the initial quality control, the rewritten multiBamSummary functionality was combined with both plotPCA and plotCorrelation functionality (Figure 1) revealing a clear separation between active histone marks (H3K4me1, H3K4me3, H3K27ac) and repressive histone marks (H3K9me3 and H3K27me3), as the largest source of variability on the sample level (Supplemental figure 3 A,B). Samples from KO and WT mNPCs separated on the PCA plot for all marks as well (Supplemental figure 3A). Both observations were corroborated by calculating the pair-wise sample correlations (Supplemental figure 3C). Finally, all ChIP samples showed a clear deviation from the diagonal in a fingerprint plot, indicating specific enrichment in certain genomic regions, with a clearer, more focal enrichment for the narrow, active marks when compared to the broad, repressive marks (Supplemental figure 3D). After peak calling for the different epigenomic marks, the transcriptional changes were linked to the epigenomic marks by correlating the estimated expression changes with the epigenomic signal changes for the genes of interest (Figure 4B). Subsequent aggregation plots revealed a distinct shift in signal intensity across the seven layers between KO and WT cells (Figure 4C). Furthermore, the differences are clearly consistent across replicates, and hold true at the single-locus level, indicating that the identified regions are biologically relevant markers of the MSL2-mediated epigenomic landscape in murine NPCs (Figure 4D).

**Figure 4.**
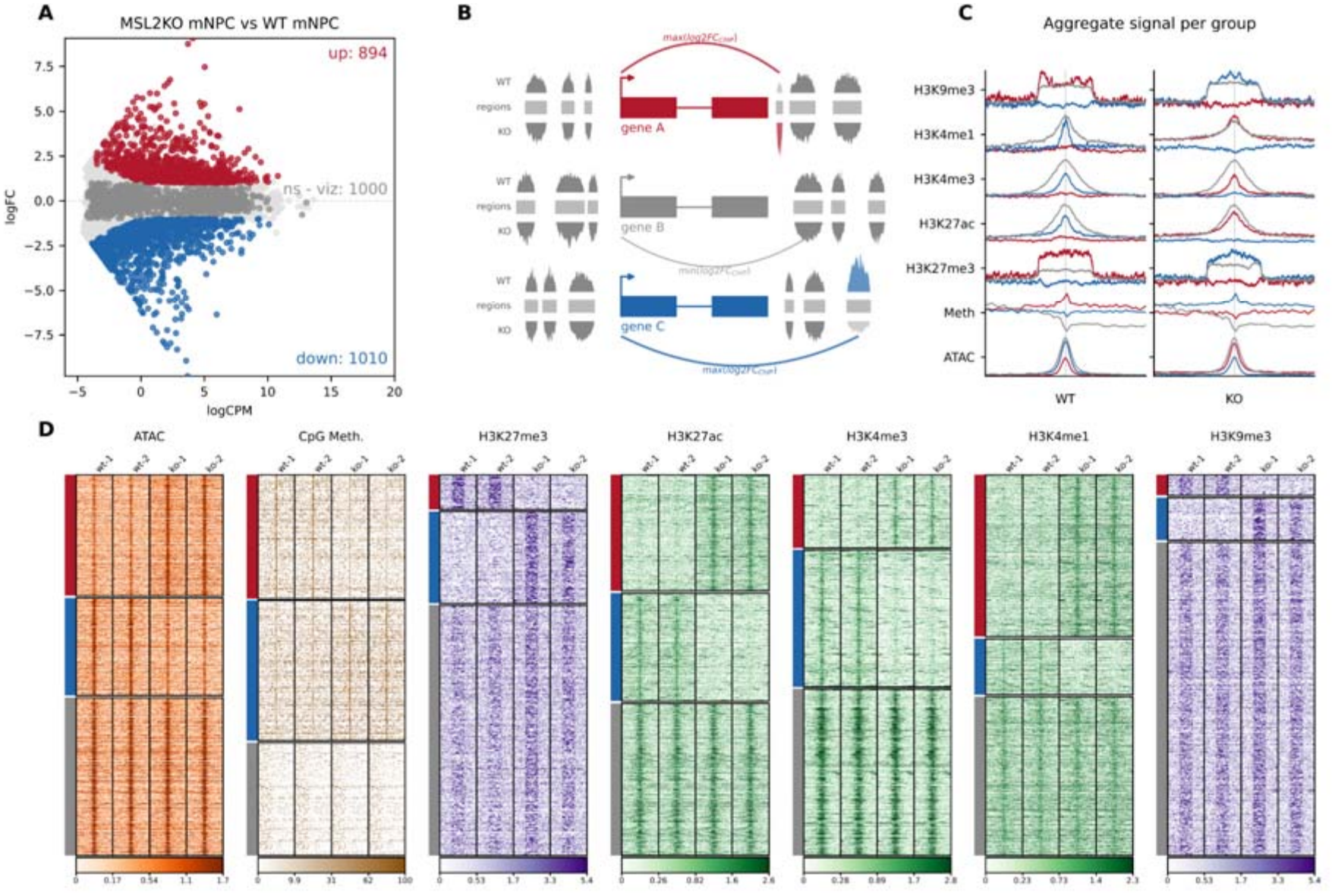
Usage of deepTools on a complex epigenomic dataset. (A) Shown are differentially expressed genes between MSL2 KO mNPCs and WT mNPCs. Upregulated (in MSL2 KO) genes are highlighted in red, downregulated (in MSL2 KO) genes are highlighted in blue. The 1,000 randomly selected non-differential genes are highlighted in dark gray. (B) Schematic overview of the selection of regions associated with the genes of interest. Per gene of interest, the signal log2 fold changes for the epigenomic marks were calculated and those with the largest difference were retained for visualization. (C) For the selected regions, aggregates were calculated and visualized using the deepTools plotProfile functionality. (D) The individual signals were visualized using deepTools’ plotHeatmap. Throughout the figure, the annotation colors (red, blue and gray) correspond to the regions associated with the up, down, and selected non-differential genes, respectively. ATAC signal is denoted in orange, DNA methylation signal is denoted in brown, repressive histone marks (H3K27me3 and H3K9me3) are denoted in purple and active histone marks (H3K27ac, H3K4me1 and H3K4me3) are denoted in green.

### Galaxy integration

Beyond the computational benchmarks and biological utility, we ensure that deepTools4 remains accessible to the scientific community by updating the Galaxy wrappers to the latest version, carefully incorporating the newly introduced parameters while removing obsolete ones. As part of the Galaxy integration, we also built a Conda package and generated a Docker container available through the BioContainers registry^19^, thereby making the software accessible to the broader bioinformatics community. The deepTools suite is widely used in Galaxy Training Network^20^ tutorials covering a broad array of omics analyses. Common use cases include assessing immunoprecipitation efficiency, generating input-normalized coverage tracks, and visualizing coverage over regions of interest using heatmaps. We updated four tutorials pertaining to ATAC-seq analysis (https://gxy.io/GTN:T00137), CUT&Run analysis (https://gxy.io/GTN:T00424), ChIP-seq enrichment analysis (https://gxy.io/GTN:T00140) and ChIP-seq peak calling analysis (https://gxy.io/GTN:T00143). These tutorials were modernized and corresponding answer histories for each tutorial were updated. In addition, we updated the associated Galaxy workflows, added missing metadata, and incorporated workflow tests in accordance with Galaxy workflow maintenance best practices (Figure 5). All updated Galaxy workflows are available through WorkflowHub under the “epigenetics” tag.

**Figure 5.**
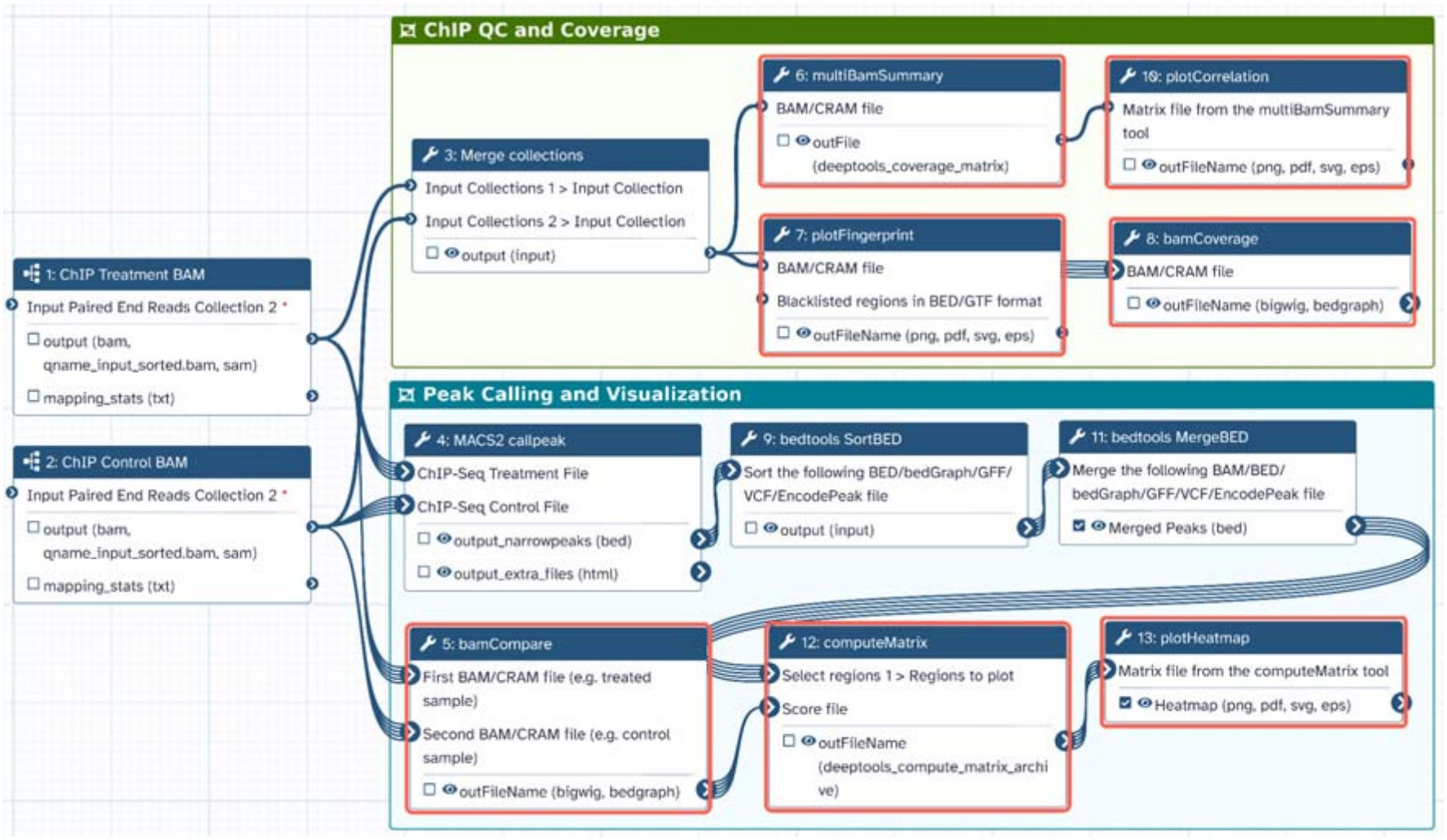
Usage of deepTools in a typical Galaxy workflow. The workflow illustrated in this figure is adapted from the ChIP-seq tutorial https://gxy.io/GTN:T00140. The workflow uses deepTools for ChIP quality control, normalization as well as downstream results interpretation. The tools used in the workflow are highlighted in red.

## Discussion

The exponential growth of both sequencing data depth as well as number of samples has created a significant “analysis bottleneck” in epigenomics. While tools for primary processing have scaled efficiently, the iterative process of quality control, normalization, and visualization has historically remained a fragmented and computationally expensive stage of the pipeline. With the release of deepTools4, we address these challenges by transitioning the suite’s core computational engine to Rust. This provides a framework that is not only significantly faster but also more robust and scalable for the next generation of genomic datasets. The transition was accompanied by a shift toward a more modular architecture and the implementation of an extensive, multi-layered testing framework. By decoupling the high-performance Rust backend from the flexible Python CLI, we have created a sustainable development pipeline. This modularity, backed by rigorous automated testing, paves the way for the seamless integration of new tools and functionalities, ensuring that deepTools4 can evolve rapidly as new sequencing modalities emerge. The most immediate impact of the update is the dramatic reduction in computational overhead. Our benchmarking across different genomes, including the exceptionally large and complex hexaploid wheat genome, demonstrates that the Rust-based backend provides consistent speed-ups across a wide spectrum of computational resources. This performance gain is particularly critical for the ‘computeMatrix’ functionality, which is often the most time-consuming step in epigenomic workflows. By reducing the time required for signal aggregation, deepTools4 transforms the analysis from a batch-processing task into a more interactive exploration. Researchers can now iterate through different normalization strategies and filtering thresholds in a fraction of the time, facilitating a more nuanced and thorough interpretation of biological data. Beyond performance, the sustainability of bioinformatics software is a growing concern. By implementing a three-tiered testing framework (integrating Rust unit tests, Pytest, and Planemo^21^), we have ensured that the core functionalities of deepTools4 are rigorously validated across different environments at different levels. Combined with the integration into the Galaxy platform, deepTools is ensured to remain accessible to a broad spectrum of users. The utility of this modernized framework is exemplified by our multimodal analysis of MSL2-mediated regulation in murine NPCs. The ability to integrate RNA-seq with seven distinct epigenomic layers and visualize these complex relationships demonstrates that deepTools4 can handle high-dimensional data without sacrificing clarity or performance. Looking forward, as the field moves toward single-cell epigenomics and long-read sequencing technologies, the demand for high-performance tooling will only increase. The architectural shift to Rust positions deepTools4 to adapt to these modalities. Future developments will focus on expanding the suite’s ability to handle these datatypes. In conclusion, deepTools4 provides a scalable, reproducible, and high-performance solution that bridges the gap between raw sequencing reads and biological discovery, ensuring that the capacity to analyze genomic data keeps pace with the capacity to generate it.

## Methods

### Implementation

To start the deepTools4 re-implementation, performance-critical components were identified as those parsing and filtering alignment files (BAM/CRAM) and aggregating and combining parsed alignment files over bins and regions. The re-implementation of those was done in Rust and exposed to Python via PyO3 bindings^22^ as a single library using maturin (https://github.com/pyo3/maturin). Input and output file processing relies on rust-htslib^7^ for BAM/CRAM files, and bigtools^8^ for bigWig data formats. This significantly reduces per-read processing overhead. Furthermore, the Rust rewrite replaces Python’s multiprocessing library with true shared-memory parallelism by relying on the Rayon library, meaning that threads can scale without inter-process serialization overhead. Compiled wheels are prebuilt for Linux and macOS, such that end users can install deepTools without needing a compiler.

### Benchmark

To quantify the performance impact of the rewrite, 6 publicly available datasets were identified from two model organisms (*Homo sapiens* and *Triticum aestivum*). In total three modalities were included. An RNA sequencing dataset (SRA accessions SRR28012902 and SRR27822150), a ChIP sequencing dataset (SRA accessions SRR28592124 and SRR1686799) and a WGS dataset (SRA accessions SRR15494527 and SRR27887047). Raw data was downloaded, and the FASTQ files were aligned using snakePipes (version 3.1.0) with default settings^15^. The GRCh38 reference genome was used for the *Homo sapiens* samples, while the IWGSC reference genome version 1.0 (accession number GCA_900519105.1) was used for the *Triticum aestivum* samples^23^. In brief, raw data was trimmed using Cutadapt (version 4.6)^24^ and subsequently aligned with either Bowtie2 (version 2.4.5)^25^ for the WGS and ChIP datasets, or STAR (version 2.7.10b)^26^. The resulting BAM files were converted to CRAM files using samtools (version 1.20)^27^ to be used as the starting point for the benchmark workflow. The rewritten bamCoverage, bamCompare, computeMatrix, multiBamSummary and alignmentSieve functions were compared against the latest Python counterparts (from deepTools version 3.5.6). For both model organisms, the tools were run on the identical input files across a range of threads between 1 and 20. Every setting was run five-fold on a high-performance server equipped with AMD EPYC 9954 processors. The replicates were performed to ensure that runtime deviations due to the file system were taken into account. Snakemake’s built-in benchmark directive was used to record runtime and peak memory usage (maximum resident set size). The resulting aggregation and visualization of the benchmark statistics were visualized using Matplotlib (version 3.11.1)^28^ and seaborn (version 0.13.2)^29^.

### Example data and processing

Publicly available epigenomic data was used to demonstrate deepTools4 in a realistic setting ^18^. A Snakemake workflow was set up that processes ChIP-seq, ATAC-seq, whole-genome bisulfite (WGBS) sequencing and RNA-seq data from both control and MSL2 knockout NPCs. Data pre-processing was done by downloading the raw data from SRA project PRJNA762170. Similar to the benchmark data, snakePipes was used to process the data, using Cutadapt to do adapter and quality trimming, followed by either Bowtie2 for alignment (ChIP-seq and ATAC-seq data), STAR (RNA-seq data) or bwa-meth (version 0.2.5)^30^ (WGBS data) against GRCm39. The resulting BAM files were converted to CRAM files using samtools (version 1.20) and used as a starting point for the example workflow. ChIP-seq data from five histone marks (H3K27ac, H3K4me1, H3K4me3, H3K27me3, H3K9me3) were processed using bamCompare against their matched input to generate log2 normalized bigWig files for narrow marks (H3K27ac, H3K4me1 and H3K4me3) and subtracted bigWig files for broad marks (H3K27me3, H3K9me3), after reads were extended to 150 bps, centered and blacklist regions were removed. The blacklist used was sourced from ExcludeRanges using the GRCm39 reference genome^31^. ATAC-seq coverage tracks were generated using the bamCoverage functionality, generating RPKM-normalized, smoothed tracks over a binsize of 10 basepairs, which were subsequently log-transformed using wiggleTools^32^. DNA Methylation signal was aggregated using MethylDackel (version 0.6.1) (https://github.com/dpryan79/methyldackel). Differentially expressed genes were identified from RNA data by using edgeR (version 4.8.2)^33^ on the count matrix generated using featureCounts (version 2.1.1)^34^. Only genes with a false discovery rate below 0.01 and an absolute log2FoldChange larger than 1 were considered differentially expressed. From the non-differentially expressed genes, 1,000 were selected at random to represent the ‘non-differential group’. Peaks for ChIP-seq and ATAC-seq data were called with MACS3 (version 3.0.4)^35^ using matched input as control for the ChIP marks. Broad histone marks (H3K27me3, H3K9me3) were called with ‘--broad’ setting, and ATAC-seq peaks were called with the ‘--nomodel --shift -75 --extsize 150’ settings. Replicate peak sets were combined and collapsed into a single consensus set of peaks using bedtools (version 2.31.1)^36^. Peak-to-gene annotations were done using UROPA (version 4.0.3)^37^, with an allowed peak-to-gene distance of 100kb in either direction. For CpG methylation, rather than calling discrete peaks, CpG sites within a 3kb window of the gene start site were considered for downstream analysis. For visualization, only one peak per gene was retained, with the strongest WT vs MSL2-KO signal (or weakest, in case of the non-differentially expressed gene set). Quantification of the signal over the regions was performed using deepTools4’s multiBamSummary functionality. Subsequent aggregate profile plots and heatmaps were generated by using deepTools4’s computeMatrix, plotProfile and plotHeatmap functionality.

## Resource availability

deepTools4 is available as open-source software and can be accessed via GitHub (https://github.com/deeptools/deepTools/). For ease of installation, the suite is distributed through both PyPI (https://pypi.org/project/deeptools/) and Bioconda (https://bioconda.github.io/recipes/deeptools/README.html). Comprehensive user documentation, tutorials, and API references are hosted via ReadTheDocs (https://deeptools.readthedocs.io/en/latest/). Additionally, deepTools4 is fully integrated on the European Galaxy platform (https://usegalaxy.eu/tools/list?tag=deepTools), making its high-performance capabilities accessible via a graphical user interface and available to install on any Galaxy instance via the Galaxy toolshed (https://toolshed.g2.bx.psu.edu/view/bgruening/suite_deeptools). To ensure reproducibility, the complete Snakemake workflows used for the performance benchmarking and the multimodal biological examples presented in this study are available in a dedicated repository (https://github.com/deeptools/deeptools4_benchmark). All raw data used for both the benchmark and example workflow are available online via their accession numbers, the processed data used in the example and benchmark setting can be accessed via Zenodo accession number 21529137 (https://zenodo.org/records/21529137). Both Snakemake workflows download the required processed files directly from Zenodo.

## Acknowledgements

We would like to thank all users of deepTools for their valuable feedback, bug reports and suggestions. We acknowledge project support from the Max Planck Computing and Data Facility and the Scientific Data Processing Unit from the Max Planck Institute of Immunobiology and Epigenetics, in particular support from Wolfgang Burger and Christian Pagel. This work was supported by funds provided by the Max Planck Society. We acknowledge support for the European Galaxy project from the German Federal Ministry of Research, Technology and Space (BMFTR) grant 031 A538A, de.NBI-RBC and the Ministry of Science, Research and the Arts Baden-Württemberg (MWK) within the framework of LIBIS/de.NBI Freiburg. T.M. acknowledges funding from the German Research Foundation (DFG RTG2344 *MeInBio*).

## Author contributions

WD conceptualized the study, wrote the backend software, wrote the unit-tests, supported the benchmark effort, analyzed the example dataset and wrote the manuscript. AS wrote the backend software and supported the benchmark effort. WM wrote the backend software, supported the Galaxy integration, and ensured Bioconda availability. MN updated the visualization part, wrote the unit-tests, and analyzed the example dataset. GE wrote the unit-tests and initiated and supported the benchmark effort. PV wrote the unit-tests and supported the Galaxy integration. SM wrote the unit-tests and supported the Galaxy integration. JC updated the visualization part. LR wrote the backend software. LLD wrote the backend software. BG conceptualized and supervised the study and supported the Galaxy integration and Bioconda integration. TM conceptualized and supervised the study.

## Declaration of generative AI and AI-assisted technologies

The authors declare that no generative AI was used for the generation of images, figures or artwork. The authors declare that generative AI tools were used to improve the language and readability of the manuscript, namely gemma4 model 31B-it. All authors reviewed and edited the suggested content and take full responsibility for the content in the manuscript. Additionally, AI tools were used to review, optimize and suggest code changes, namely Qwen models 3.6-27B and 3.8-27B and openAI’s GPT-3.5. All proposed changes and feedback were reviewed and verified by one of the authors before they were incorporated.

