## Supplemental Tables and Figures for "deepTools4: Faster, Scalable, and Modernized Genomic Data Analysis"

### Supplemental Information

**Supplemental table 1.** Sample information for the publicly available data used in the benchmark.

| **Organism** | **Modality** | **total reads (M)** | **% aligned** | **% dups** | **avg. read length** | **mean insert size** | **% genome covered** | **average depth** |
| --- | --- | --- | --- | --- | --- | --- | --- | --- |
| *Homo sapiens* | ChIP | 33.4 | 95.63 | 0.0 | 149 | 303.2 | 57.8 | 1.53 |
| *Homo sapiens* | RNA | 251.4 | 99.47 | 0.0 | 101 | 978.5 | 9.1 | 3.08 |
| *Homo sapiens* | WGS | 966.2 | 98.33 | 12.1 | 101 | 408.8 | 93.8 | 27.14 |
| *Triticum Aestivum* | ChIP | 64.1 | 77.93 | 0.0 | 101 | 146.3 | 18.9 | 0.35 |
| *Triticum Aestivum* | RNA | 25.7 | 88.29 | 0.0 | 151 | 394.6 | 1.0 | 0.19 |
| *Triticum Aestivum* | WGS | 1092.6 | 97.75 | 0.0 | 150 | 337.0 | 94.4 | 11.01 |

**Supplemental table 2**. Overview of the hierarchical testing framework for the deepTools suite. The framework consists of three distinct levels designed to ensure robustness, reproducibility, and ease of future development.

|  | **Test framework** | **Language** | **Target** | **Level** | **Number of tests** |
| --- | --- | --- | --- | --- | --- |
| 1 | Rust unit tests | Rust | core functionality | Module | 48 |
| 2 | Pytest | Python | CLI  visualization metrics  statistics | Tool | 232 |
| 3 | Planemo | Rust, Python | Galaxy integration | Workflow | 54 |

**Supplemental table 3.** Sample information for the data used in the example workflow.

| **Sample** | **total reads (M)** | **avg. read length** | **insert size mean** | **insert size sd** |
| --- | --- | --- | --- | --- |
| WT_H3K27ac_rep1 | 15.8 | 88.0 | 269.1 | 147.4 |
| WT_H3K27ac_rep2 | 15.9 | 88.0 | 273.3 | 147.7 |
| WT_H3K27me3_rep1 | 32.2 | 88.0 | 294.6 | 171.5 |
| WT_H3K27me3_rep2 | 33.1 | 88.0 | 299.1 | 171.3 |
| WT_H3K4me1_rep1 | 16.1 | 88.0 | 267.2 | 151.2 |
| WT_H3K4me1_rep2 | 17.1 | 88.0 | 271.3 | 151.1 |
| WT_H3K4me3_rep1 | 13.9 | 88.0 | 239.4 | 140.1 |
| WT_H3K4me3_rep2 | 14.5 | 88.0 | 250.0 | 141.5 |
| WT_H3K9me3_rep1 | 34.6 | 88.0 | 318.5 | 178.1 |
| WT_H3K9me3_rep2 | 35.5 | 88.0 | 320.9 | 177.8 |
| WT_input_rep1 | 76.5 | 101.0 | 232.5 | 108.5 |
| WT_input_rep2 | 78.6 | 101.0 | 247.8 | 119.2 |
| WT_RNA_rep1 | 28.7 | 101.0 | 952.7 | 1816.4 |
| WT_RNA_rep2 | 28.5 | 101.0 | 975.2 | 1835.9 |
| WT_ATAC_rep1 | 76.6 | 101.0 | 212.6 | 103.4 |
| WT_ATAC_rep2 | 82.7 | 101.0 | 216.0 | 108.5 |
| WT_BS_rep1 | 63.8 | 101.0 | 262.8 | 124.5 |
| WT_BS_rep2 | 58.9 | 101.0 | 258.2 | 116.4 |
| MSL2-KO_ATAC_rep1 | 89.0 | 101.0 | 223.4 | 117.2 |
| MSL2-KO_ATAC_rep2 | 81.8 | 101.0 | 222.8 | 114.6 |
| MSL2-KO_BS_rep1 | 811.9 | 101.0 | 256.1 | 117.5 |
| MSL2-KO_BS_rep2 | 601.1 | 101.0 | 253.6 | 116.3 |
| MSL2-KO_H3K27ac_rep1 | 16.0 | 90.0 | 291.2 | 118.2 |
| MSL2-KO_H3K27ac_rep2 | 15.0 | 90.0 | 280.2 | 112.4 |
| MSL2-KO_H3K27me3_rep1 | 29.6 | 90.0 | 320.7 | 145.8 |
| MSL2-KO_H3K27me3_rep2 | 26.5 | 90.0 | 306.9 | 139.6 |
| MSL2-KO_H3K4me1_rep1 | 24.4 | 90.0 | 305.8 | 125.5 |
| MSL2-KO_H3K4me1_rep2 | 23.1 | 90.0 | 293.5 | 119.6 |
| MSL2-KO_H3K4me3_rep1 | 9.6 | 90.0 | 280.2 | 115.5 |
| MSL2-KO_H3K4me3_rep2 | 10.1 | 90.0 | 268.7 | 109.1 |
| MSL2-KO_H3K9me3_rep1 | 32.0 | 90.0 | 330.5 | 154.2 |
| MSL2-KO_H3K9me3_rep2 | 28.1 | 90.0 | 315.8 | 147.5 |
| MSL2-KO_input1_rep1 | 99.8 | 101.0 | 245.7 | 118.0 |
| MSL2-KO_input1_rep2 | 61.6 | 101.0 | 236.6 | 111.2 |
| MSL2-KO_RNA_rep1 | 36.2 | 101.0 | 927.6 | 1778.0 |
| MSL2-KO_RNA_rep2 | 27.9 | 101.0 | 992.0 | 1861.2 |

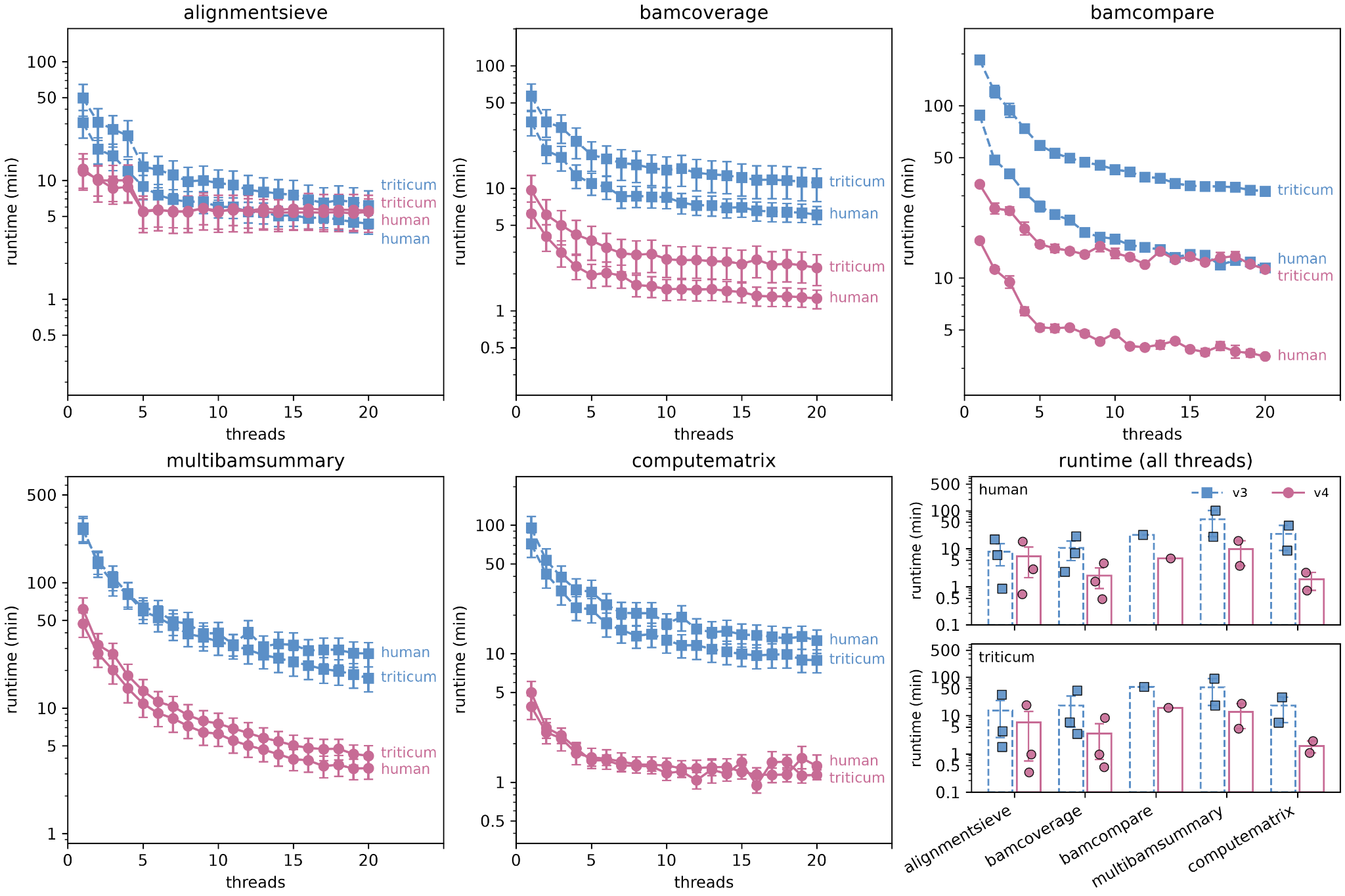

**Supplemental figure 1. Actual wall times for rewritten backend.**  Shown are the absolute runtimes (in minutes) for deepTools4 (circles, purple) compared to deepTools3 (squares, blue). Error bars denote the standard error of the mean calculated from five independent runs. The last panel depicts the average runtime (in minutes) split up per organism.

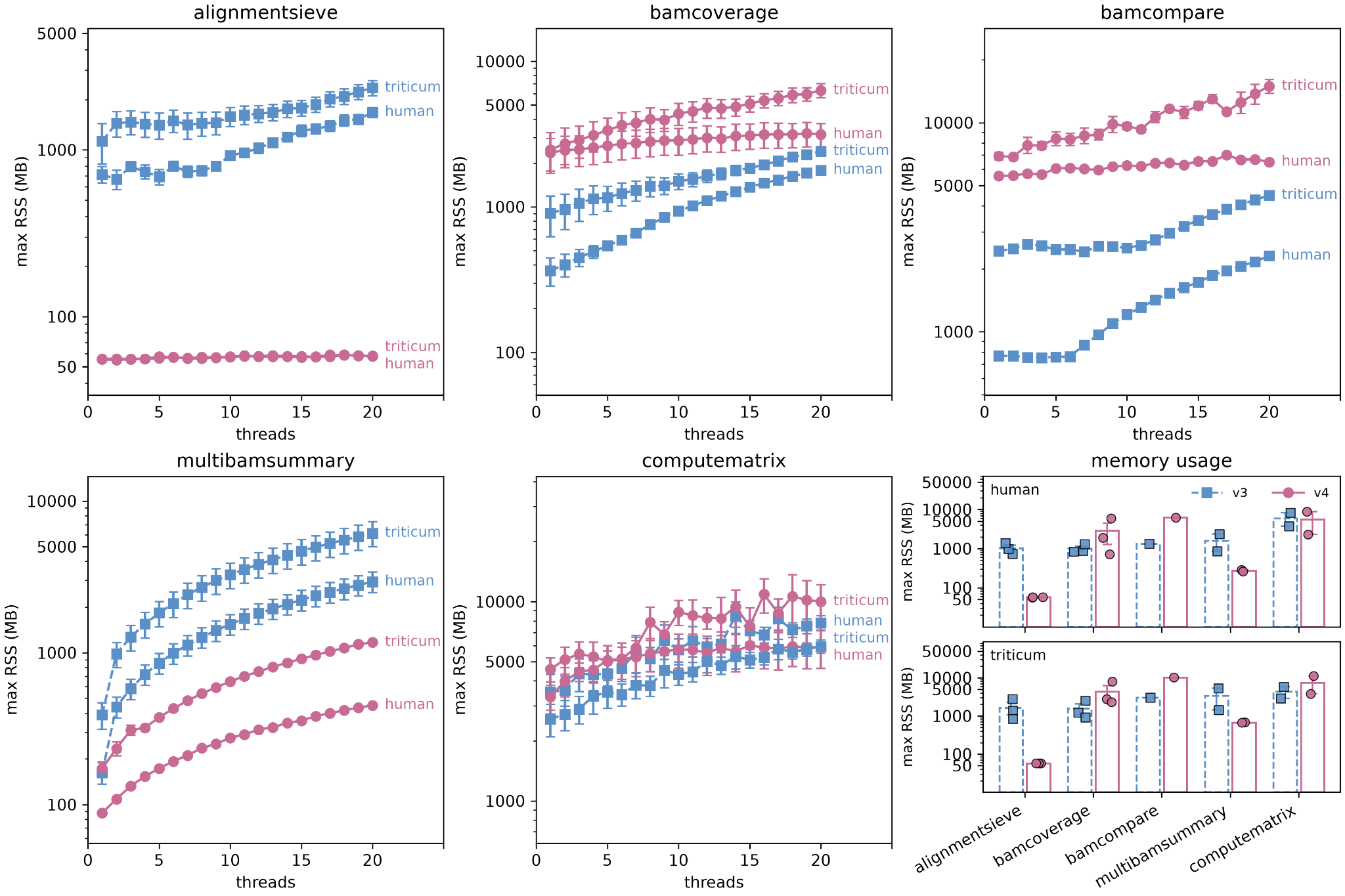

**Supplemental figure 2. Memory usage for rewritten backend.**  Shown is the maximum resident set size (in MB) for deepTools4 (circles, purple) compared to deepTools3 (squares, blue). Error bars denote the standard error of the mean from five independent runs. The last panel depicts the average maximum resident set size (MB) split up per organism.

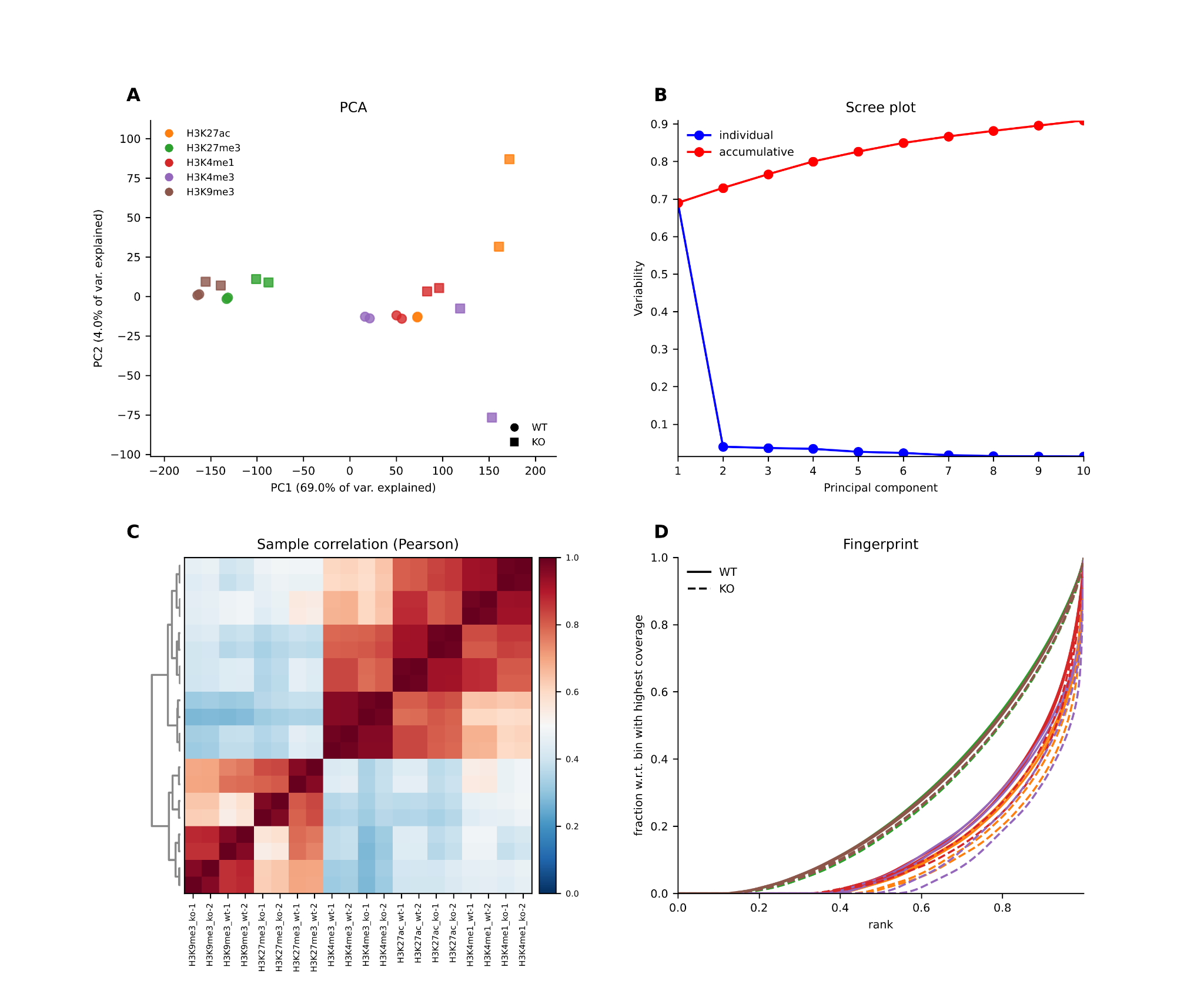
**Supplemental figure 3**. **Visualization examples in the context of quality control for the publicly available ChIP samples.** (A) Shown is the PCA plot based on log2-transformed counts on 10kb genomic bins. Different colors denote different histone marks, while circles denote WT samples and squares denote KO samples. (B) Scree plot associated with the PCA plot denoting the individual fraction of variance explained (blue) or the cumulative variance explained (red) per additional principal component. (C) Heatmap showing pair-wise Pearson correlations between samples calculated using the same underlying data as the PCA plot. (D) Genomic bins (ranked by increasing read coverage) versus the cumulative fraction of total reads. Individual curves denote a single sample, following the same color encoding as in the PCA plot. WT samples are denoted as solid curves, while KO samples are denoted as dashed curves. A diagonal line would denote uniform coverage across the genome.
